# The macroevolutionary dynamics of mammalian sexual size dimorphism

**DOI:** 10.1101/2023.05.31.543075

**Authors:** Megan E. Jones, Catherine Sheard

## Abstract

Sexual size dimorphism (SSD) is a common phenomenon across the animal kingdom. Mammals are unusual in primarily displaying male-biased SSD, where males of a species are typically larger than females. The driving factors behind the evolution of this SSD have been much debated, with popular hypotheses including the influence of mating system and social organisation via sexual selection, dietary niche divergence, and the role of body size (Rensch’s rule). Here, we investigate the macroevolutionary origins and maintenance of SSD among mammals, using phylogenetic general mixed linear models and a comprehensive global dataset to evaluate correlations of diet, body mass, seasonality, social organisation, and mating system with SSD type. We find that SSD as a whole is lost at a greater rate than it is gained, with female-biased SSD being particularly unstable. Non-monogamy, vertebrate prey consumption, and temperature seasonality correlate with male-biased SSD, while polyandry correlates with female-biased SSD, and both types of SSD are positively correlated with body mass. This is in partial contrast to the predictions of Rensch’s rule, which predicts that female-biased SSD would correlate negatively with body size. Taken together, our results highlight the importance of considering multiple ecological and social drivers when evaluating the macroevolutionary trajectory of sex differences in body size.

## Introduction

Sexual size dimorphism (SSD), the phenomenon where members of one sex of a species are consistently and significantly larger than the other, can be found throughout the animal kingdom. In most animals, including invertebrates and ectothermic vertebrates, SSD tends to be female-biased – that is, with larger females than males [1]. Mammals are thus unusual, as SSD among mammals is most often male-biased [1, 2], with female-biased SSD only predominant among a few orders, such as the Chiroptera (bats) and Lagomorpha (rabbits, hares and relatives) [3]. The evolution of SSD among mammals has been the subject of extensive study at various scales (e.g. [1, 3, 4]), but patterns of interspecific variation in SSD across the class have rarely been examined.

Perhaps the most commonly cited potential driver of interspecific variation in mammalian SSD is sexual selection, specifically as quantified by differences in mating system [1, 2, 4, 5]. In species with male-biased SSD, larger males are generally thought to achieve greater reproductive success, as they have advantage over smaller males in direct competition for mates (including physical access and/or female choice) [3, 5, 6]. In particular, in polygynous species, a single male will monopolise multiple females. This means that, in theory, any competitive advantage between males (such as body size) will have a greater effect on their lifetime reproductive success than it would in monogamous species, as the variance in the number of females any given male might mate with is much higher. However, recent genetic paternity studies, among others, have found that the biological reality is more complicated [5, 7-9], with variance in male reproductive success not necessarily increasing in polygynous species; thus, a broad-scale link between mating system and SSD may not be as straightforward as previously thought. This potential species-level relationship between mating system and SSD is further mediated by social organisation, or sociality. Both intrasexual competition and the opportunities for mate choice are thought to be greater among group-living species than in solitary ones [10, 11], suggesting that selective pressure for dimorphism might be stronger among these more social species.

Interspecific variation in SSD has also been linked to body size and allometry. Rensch’s rule proposes that, among species with male-biased SSD, the degree of dimorphism will increase with increasing body size; meanwhile, female-biased SSD will decrease with increasing body size [12-15]. Rensch’s rule seems to hold true commonly – although far from universally –across mammals with male-biased SSD; however, those species with female-biased SSD very rarely follow this pattern [3, 12, 16]. Importantly, Rensch’s rule does not itself suggest a mechanism by which increasing size will lead to more pronounced male-biased SSD. Instead, various additional hypotheses have subsequently been proposed to explain this, with many linking back to the sexual selection hypothesis discussed previously (e.g. [12, 13]). For example, Sibly *et al*. [15] observe that mammal groups which follow Rensch’s rule – such as Bovidae, Cervidae, and Macropodidae – tend to also increase group size with body mass. The authors link this back to sexual selection, using group size as a proxy for polygyny, which in turn is considered a proxy for sexual selection intensity. Rensch’s rule has also been linked to fecundity selection. Unlike in many animal groups, fecundity decreases and reproduction is more energetically costly in females of larger mammal species [3, 17]; this may lead to female body size being more constrained than male body size in these larger species [18].

Another hypothesis proposes that the evolution of SSD is driven by dietary niche divergence [1, 8, 19, 20]. Males and females of different sizes may occupy different dietary niches, and thus high SSD may decrease levels of intraspecific competition for food. Thus, species with more specialised diets may face greater pressure for either male- or female-biased SSD, though note that, in isolation, this explanation is less able to explain why specifically male-biased SSD would be predominant among mammals. Evidence for this hypothesis at the macroevolutionary scale is scarce, though a version of this may explain the observation that species in the order Carnivora, which primarily feed on terrestrial vertebrates, tend to show more pronounced male-biased SSD [17, 21, 22]. The uneven distribution of terrestrial vertebrate prey potentially leads to more intense intraspecific competition among species that predate on these animals, in comparison to species which rely on more evenly distributed food sources, such as herbivores or insectivores [17, 21].

The dietary niche divergence hypothesis is difficult to investigate at the macroevolutionary level, not only due to the challenges in quantifying dietary niche at this scale [23, 24], but also due to the difficulty in distinguishing cause and effect. It is entirely plausible that SSD could initially evolve through other means, with dimorphism itself then driving a subsequent separation of the dietary niches [1]. In this case, however, dietary niche partitioning would still be a factor in maintaining SSD across evolutionary time. Moreover, this is not the only hypothesis where cause and effect may be difficult to disentangle, a major limitation of a comparative approach. Broad studies, however, of species across many ecological contexts, could begin to pinpoint which of these taxon-specific patterns may be the consequence of general evolutionary principles, and which might instead be the by-product of more local or particular processes.

A final major hypothesis for the drivers of mammalian SSD at wide-ranging scales is a potential link between dimorphism and climate variability (seasonality) [4]. The studies supporting this correlation have generally been limited to specific taxa, and present conflicting evidence of the effects of increased seasonality. For example, Kappeler *et al*. [25] and Kappeler [26] suggest that high seasonality may reduce SSD among prosimian primates, while Garel *et al*. [27] find that moose populations show increased SSD under more seasonal conditions. Such an inter-specific relationship between seasonality and SSD, however, might be complicated by co-varying relationships between both variables and social organisation [28], further underscoring the need for analytic approaches within a multivariate framework.

Most of the existing literature on SSD evolution among mammals investigates a limited range of taxa – such as a single order, family, or species – and many fail to consider the additive effects of multiple hypotheses. None of the five hypotheses here are mutually exclusive, and indeed, many are explicitly related to one another. To address these complex relationships at the global, class-level scale, we here first use a dataset of SSD in 5,261 species of mammals and describe the macroevolutionary transitions among male-biased SSD, female-biased SSD, and monomorphism. Then, on a more restricted dataset of 1,662 species from 134 families, we use Bayesian phylogenetic mixed models to investigate how body size, mating system, social organisation, dietary niche, and seasonality correlate with these transitions on a global scale.

## Methods

### Data collection

Sexual size dimorphism (SSD) is here classified as male-biased, female-biased, or monomorphic and was collected for 5,949 species based on the text descriptions in the *Handbook of the Mammals of the World* (HMW) [29]. A species was classified as dimorphic if (1) the textual description explicitly stated the presence and direction of SSD; or if the following measures were listed separately by sex with one larger than the other (or, if only averages were listed, if the averages differed by at least 2%, indexed to the female): (2) body mass, (3) head-body length, (4) shoulder length, or (5) for the order Chiroptera, forearm length. Though 2% is a low threshold for the rare cases when only averages were listed, we considered the separate listing of male and female body masses by the expert HMW authors to be a strong indication that biologically meaningful dimorphism was present. These scores were recorded as a ternary variable rather than collected as an index of the degree of dimorphism, both to keep the sample size as large as possible (as easily-collated, standardised, sex-specific morphological measurements do not yet exist for all mammals) and to harness the power of phylogenetic comparative methods that measure transition rates between categorical variables.

Mass and dietary data were obtained from the PHYLACINE database [30]. To be broadly comparable across the entire class, dietary data is here considered as one of three integer scores – plants, vertebrates, and invertebrates – totalling 100 and representing the approximate proportion of each category within that species’ diet. The underlying PHYLACINE dietary dataset is primarily based on EltonTraits 1.0 [31] and MammalDIET [23], which should be consulted for further information about their data collection efforts and phylogenetic imputation strategies. Following the suggestions of Law and Mehta [21] and Law [17], we consider in particular the percentage of vertebrate prey in a species’ diet as a proxy for the potential for natural selection in favour of dietary niche divergence, in order to investigate the hypothesis that dietary niche divergence drives the evolution of SSD. This is an imperfect proxy, but comprehensive data on dietary niche divergence across mammal species is not yet available.

The mating systems of 1,874 of the mammal species were classified as monogamous, polyandrous, polygynous, polygynandrous (termed “promiscuous” by many sources, though see [32]), or mixed (i.e. showing multiple types of mating system). These data were gathered from the *Handbook of the Mammals of the World* [29], the Animal Diversity Web [33], and the Encyclopedia of Life [34]; further information can be found in the supplementary material. Although observed mating system is likely a poor proxy for the actual degree of sexual selection occurring within a species, this trait is favoured by comparative studies because it can be measured across such a large number of species.

Social organisation data was also obtained where possible from the *Handbook of the Mammals of the World* [29], the Animal Diversity Web [33], and the Encyclopedia of Life [34], scored as “solitary”, “pair”, and “group”. Where intraspecific or temporal variation was indicated, the more social score was used (e.g., group > pair > solitary). This information was unavailable for many species, and we suspect that the “pair” category in particular might be under-reported within our dataset.

Finally, two common proxies for climate seasonality (the average variation in climatic conditions over the course of a year) were calculated by intersecting two BioClim 2.1 [35] variables – temperature annual range (maximum temperature of warmest month minus minimum temperature of coldest month, BIO7) and precipitation seasonality (coefficient of variation, BIO15), both aggregated at the scale of 10 minutes – with the PHYLACINE range maps [30], and taking the mean value of each variable for each species. Thus, “temperature seasonality” here represents the difference in a location’s minimum and maximum monthly temperature, averaged across the range, while “precipitation seasonality” represents a measure of annual monthly variability of a location’s rainfall, averaged across the species range.

A set of 100 phylogenies was drawn at random from the posterior distribution of Upham *et al*. [36]; references to other quantities of phylogenetic trees were subsampled from this set.

### Phylogenetic comparative analyses

To first describe the macroevolutionary dynamics of SSD across all mammals (n = 5,261 species), we calculated the rates of transition among SSD states using the “MultiState” model in BayesTraits V4.0.0 [37]. All parameter priors were drawn from an exponential distribution with a mean of 10, and to ensure that the model accounted for phylogenetic uncertainty, the chain was forced to spend an equal amount of time on each tree. Two sets of models were run: one with 10 randomly-selected trees, run for 100,000 iterations per tree, and one with 100 randomly-selected trees, run for 10,000 iterations per tree. In both models, the first 1,000 iterations were discarded as burn-in, and chains were sampled every 1,000 iterations thereafter. Both sets of results were visually inspected to ensure proper mixing, and the results are qualitatively similar to one another. We present the results of the 100-tree model here, and the results of the 10-tree model can be found in the supplementary material.

Once the patterns of SSD state transition were described, we assessed correlations between these transitions and key social, ecological, and environmental variables using Bayesian phylogenetic mixed models in the package MCMCglmm [38], in R 4.2.0. We began with one foray into model selection. The inclusion of social organisation as a potential explanatory variable substantially dropped our sample size, increasing geographic and taxonomic biases to well-studied species. We therefore ran two models using the same dataset of 1,457 species: one including social organisation as a potential correlate and one without. We then compared the Deviance Information Criteria (DIC) of these two models to assess whether including this variable improved or reduced the fit of the model. The lower DIC was seen in the model without the social organisation data (ΔDIC = 5.7); therefore, all further analyses were run on a dataset of 1,662 species with four types of explanator variable (body mass, diet, mating system, and climate seasonality). For more information, as well as for a model selection procedure testing the fit of the seasonality data (the potential explanatory variable with the most tenuous macroevolutionary link), please see the supplementary information (Tables S1-6).

Our main model, a phylogenetic “categorical” model, which we hereafter refer to as the “separate-effects” model, considers the differences between monomorphism and each of female-biased and male-biased SSD as the response variable (Fig. 1). We also ran individual phylogenetic logistic regressions on each pairwise trait (male-vs female-biased: N = 714; monomorphic vs female-biased, N = 1,078; and monomorphic vs male-biased, N = 1,543) for ease of interpretation, and to more closely mirror the description of the macroevolutionary transitions.

**Figure 1:**
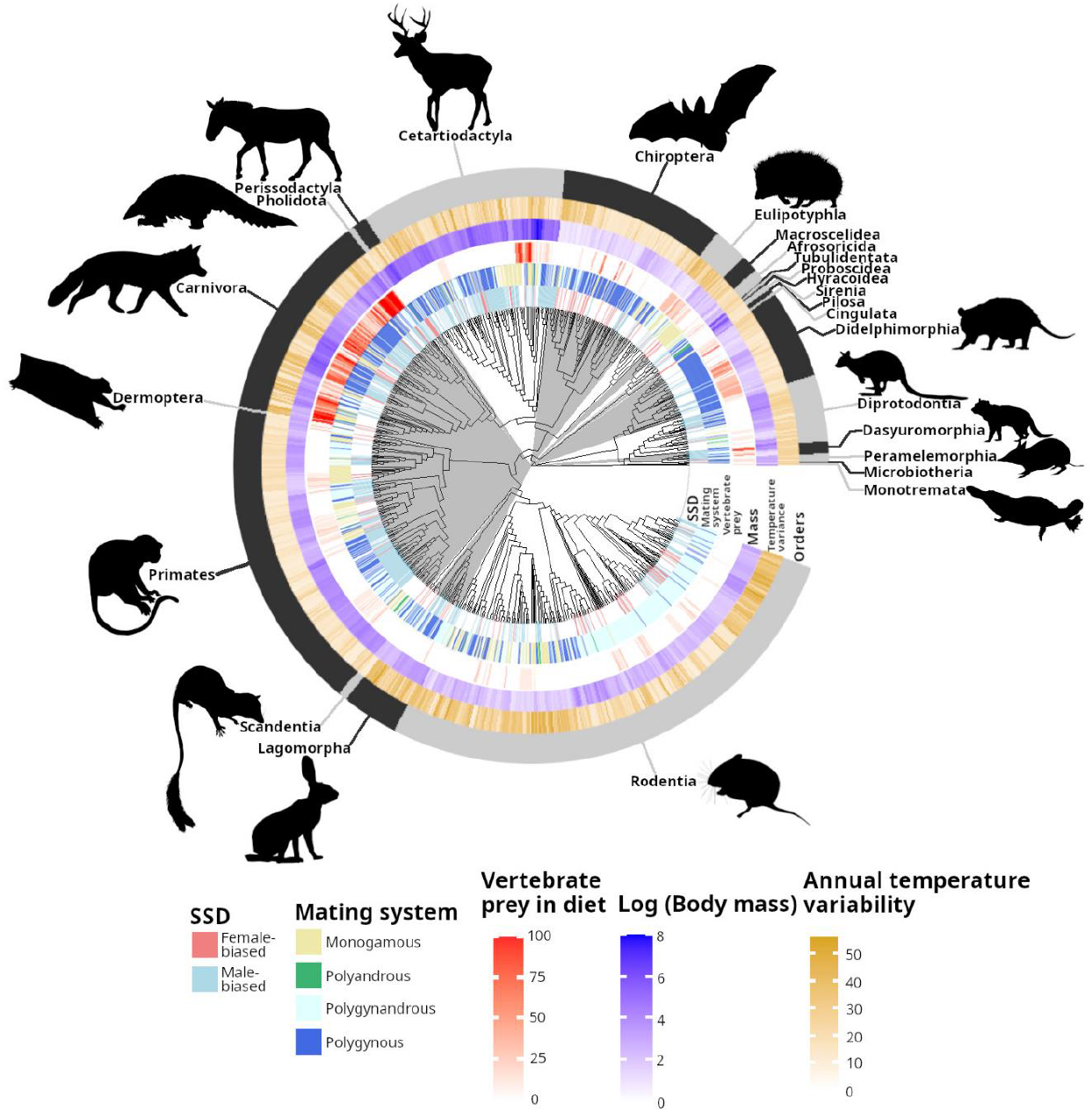
Phylogeny of Mammalia, showing types of SSD displayed by each species, along with three key traits found to significantly correlate with type of SSD (vertebrate prey in diet, mating system, and body mass). The phylogenetic topology shown is a consensus tree taken from Upham *et al*. [36]; the tree and variables were plotted in R 4.2.0, using the package ggtree [50]. Silhouettes are taken from Phylopic 2.0 and are by Tracy A. Heath, Yan Wong, Inessa Voet, Margot Michaud, Michael Scroggie, Steven Traver, T. Michael Keesey, all CC0 1.0.

We also ran two sets of sensitivity analyses on our separate-effects model. The first tested other thresholds beyond the 2% used in the macroevolutionary transition analyses; as entries listing measurement averages (rather than ranges) were relatively rare, we also calculated averages from listed measurement ranges for the purposes of these tests. We thus re-ran analyses on 1) a dataset where species with average measurements differing by < 5%, indexed to the female, were scored as monomorphic (n = 32), and 2) a dataset where species with average measurements differing by < 20%, indexed to the female, were scored as monomorphic (n = 165). The second set of sensitivity analyses considered where possible variation between body masses (the preferred measurement for our scoring system, but not always available) and linear measurements (head-body length, shoulder height, and forearm length). We thus re-ran analyses on a dataset where any species scored as dimorphic for body mass but that would not be scored as dimorphic (or would be scored as dimorphic in the other direction) for a linear measurement was considered monomorphic (n = 21). Further details on these model outputs are available in the supplementary information (Tables S7-9).

To aid coefficient comparability, all numerical variables were normalised (scaled to have a mean of 0 and variance of 1) before analysis, and mass was log-transformed. All models were run on 100 randomly-selected trees from Upham *et al*. [36]; after an initial dummy run to determine a start point, each tree was run for 55,000 iterations, with the first 5,000 discarded as burn-in, and sampling every 5,000 iterations thereafter. The residual variance was fixed to 1; a non-informative prior was selected for the phylogenetic variance (V = 1^-10^, *v* = -1); and priors for the fixed effects were kept as the default (diffuse normal priors with mean 0, variance 10^10^). The variance inflation factor (VIF, calculated based on code from [39], originally written by Austin Frank) for each fixed effect was checked and determined to be <5, and all effective sample sizes were greater than 250. Trace and density plots of the model outputs were visually examined to ensure convergence and proper mixing.

## Results

### Data overview

Of the 5,949 species in the HMW for which we were able to obtain dimorphism scores, 1,149 (19%) were recorded as having male-biased SSD and 311 (7%) as having female-biased SSD. Documented male-biased SSD was particularly prevalent in the Dasyuromorphia (carnivorous marsupials; 68 of 75, 91%), Pholidota (scaly anteaters; 6 of 8, 75%), Proboscidea (elephants; 2 of 3, 67%), Peramelemorphia (bandicoots and bilbies; 11 of 19, 58%), and Carnivora (carnivorans such as cats, dogs, bears, and weasels; 157 of 279, 56%). Documented female-biased SSD was particularly prevalent in the Macroscelidea (elephant shrews; 5 of 20, 25%), Pilosa (sloths and anteaters; 3 of 16, 19%), and Perissodactyla (odd-toed ungulates such as horses and tapirs; 2 of 16, 13%). SSD was rarely observed in many orders, including Eulipotyphla (hedgehogs and shrews; 498 of 530, 94% monomorphic), Lagomorpha (hares, rabbits, and pikas; 83 of 92, 90% monomorphic), and Rodentia (rodents; 2,138 of 2,475, 86% monomorphic). We stress, however, that many of these species potentially exhibit unrecorded SSD.

### The macroevolutionary dynamics of SSD

Transitions directly between female- and male-biased dimorphism were rare (∼21x and ∼3.4x less likely than transitions from female-biased SSD to monomorphism and from male-biased SSD to monomorphism, respectively) (Fig. 2, Table S10-11). Monomorphism was the most stable state, being ∼16x more likely to be gained from female-biased SSD than lost, and ∼4x more likely to be gained from male-biased SSD than lost.

**Figure 2:**
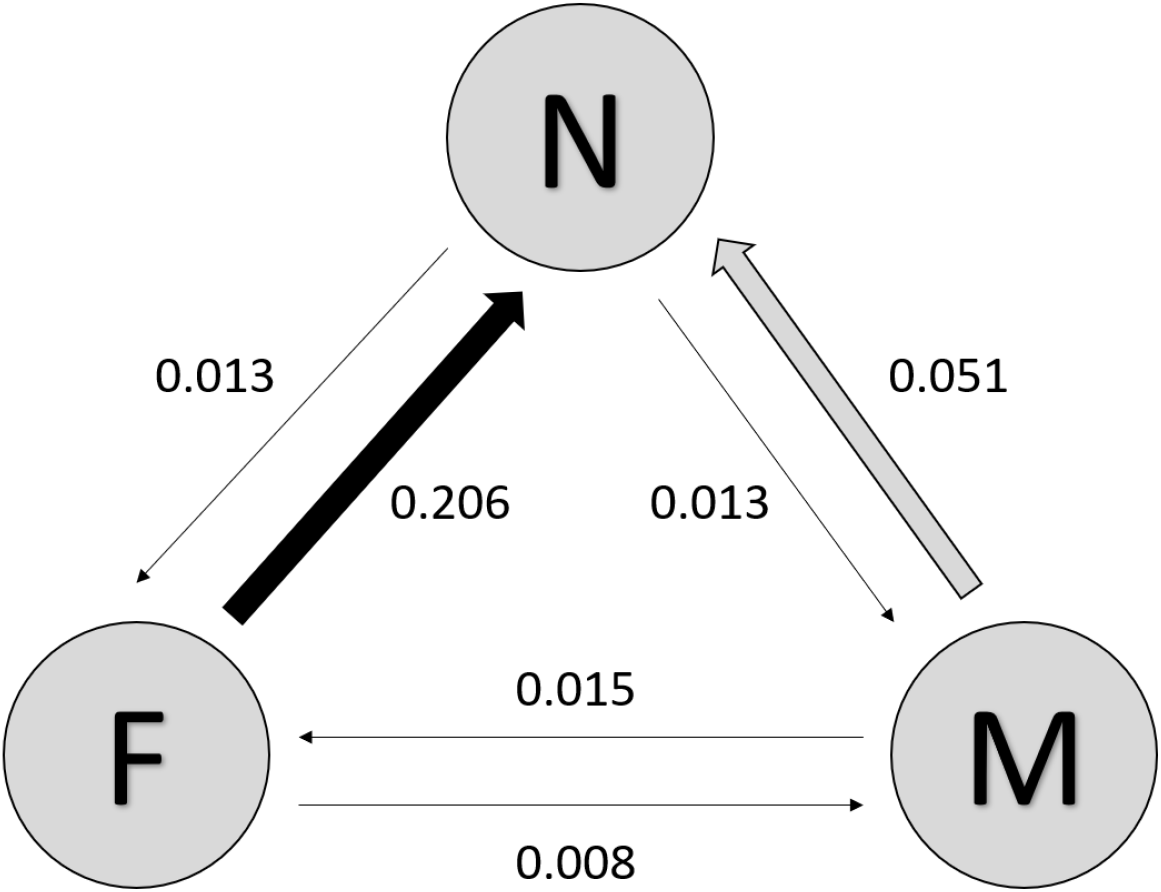
Transition rates of sexual size dimorphism (SSD). Arrows are sized and shaded to indicate transition rate values. Values are taken from a MultiState model with an equal number of iterations over 100 trees (see Materials and Methods and Table S10 for details). Key: f = female-biased SSD, m= male-biased SSD, n = no SSD.

### Social and ecological drivers of SSD

SSD is more likely in larger species (β_f_ = 1.051, pMCMC = 0.004; β_m_ = 1.462, pMCMC < 0.001; Table 1), though most of this relationship is driven by a positive correlation between body mass and male-biased SSD (Table 2). This is in partial accordance with Rensch’s rule, which both predicts that male-biased SSD should be more common in larger species and that female-biased SSD should be *less* common in larger species; we find that body mass does not significantly differentiate between female-biased SSD and monomorphism at this scale. We similarly find that male-biased SSD is related to diet (with a vertivore diet correlating with higher levels of SSD, β_m_ = 0.657, pMCMC = 0.006) and temperature seasonality (with higher levels of SSD found in regions with greater annual temperature variability, β_m_ = 0.432, pMCMC = 0.018), but no relationship between these variables and female-biased SSD.

**Table 1:** Results of phylogenetic generalised linear mixed models (MCMCglmm) for the social, ecological and environmental correlates of (m) male- and (f) female-biased SSD, versus monomorphism. Body mass is log-transformed; plants and vertebrates are dietary categories scored, along with invertebrates, out of a total of 100; mating system coefficients are against a baseline of monogamy; all continuous variables are scaled to have a mean of 0 and a variance of 1 prior to analysis. Significant correlations are highlighted in grey (pMCMC < 0.05).

|  |  |  | Coefficient | Lower 95% CI | Upper 95% CI | pMCMC |
| --- | --- | --- | --- | --- | --- | --- |
| Mass |  | f | 1.051 | 0.257 | 1.774 | 0.004 |
|  |  | m | 1.462 | 0.728 | 2.220 | <0.001 |
| Diet | Plants | f | -0.702 | -1.375 | 0.068 | 0.058 |
|  |  | m | -0.147 | -0.850 | 0.554 | 0.688 |
|  | Vertebrates | f | -0.041 | -0.589 | 0.552 | 0.866 |
|  |  | m | 0.657 | 0.183 | 1.211 | 0.006 |
| Mating system | Mixed | f | 0.239 | -0.905 | 1.354 | 0.686 |
|  |  | m | 1.058 | 0.116 | 2.057 | 0.030 |
|  | Polyandrous | f | 3.049 | 0.431 | 5.214 | 0.014 |
|  |  | m | 1.483 | -0.832 | 4.357 | 0.246 |
|  | Polygynandrous | f | 0.550 | -0.563 | 1.624 | 0.278 |
|  |  | m | 1.002 | 0.138 | 2.020 | 0.034 |
|  | Polygynous | f | -0.114 | -1.244 | 0.827 | 0.842 |
|  |  | m | 1.324 | 0.482 | 2.137 | 0.002 |
| Seasonality | Precipitation | f | -0.147 | -0.494 | 0.248 | 0.448 |
|  |  | m | -0.248 | -0.555 | 0.095 | 0.150 |
|  | Temperature | f | 0.408 | -0.007 | 0.860 | 0.056 |
|  |  | m | 0.432 | 0.043 | 0.809 | 0.018 |

**Table 2:**
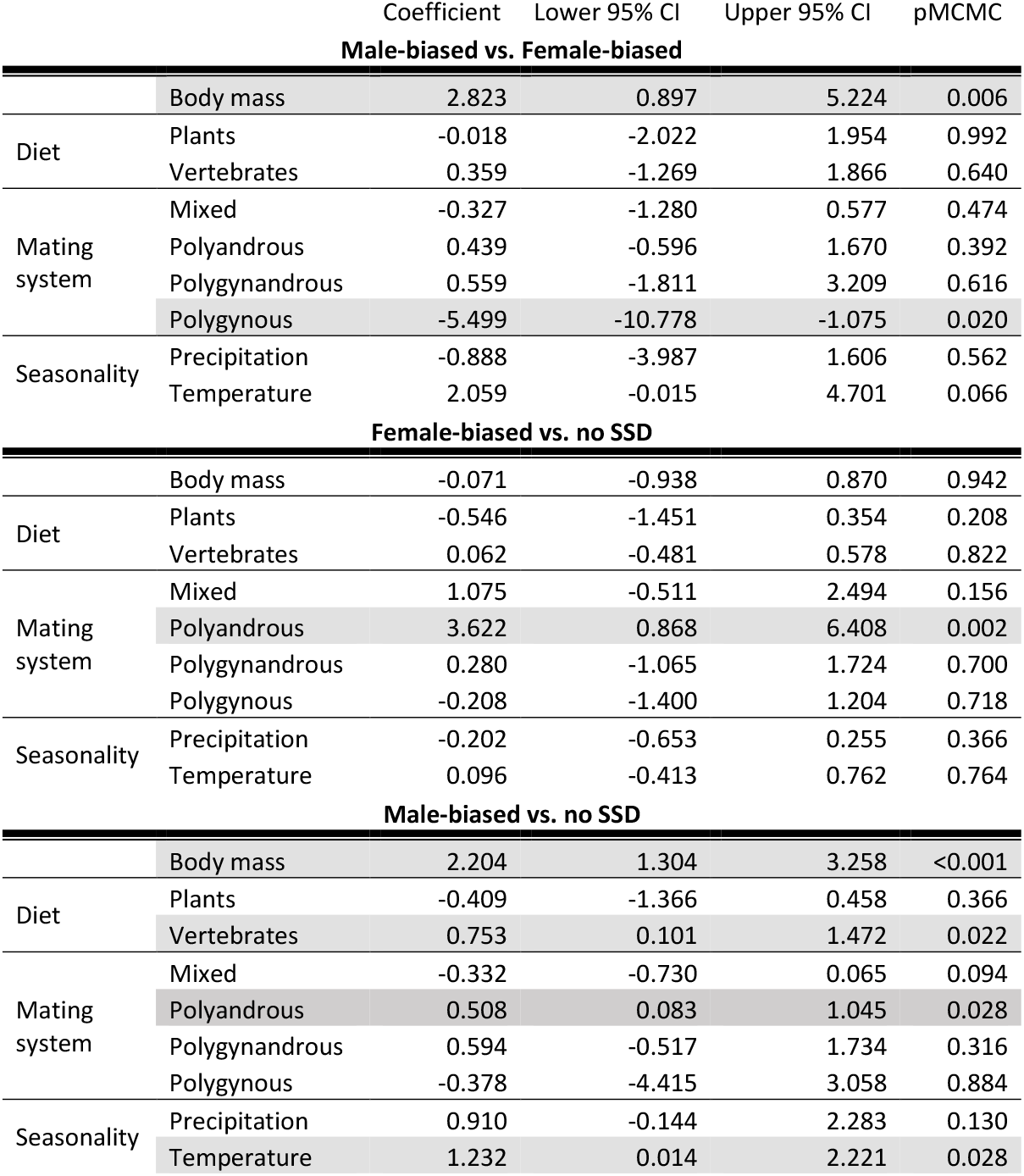
Results of phylogenetic generalised linear mixed models (MCMCglmm) for the social and ecological correlates of different types of SSD, considered as pairs (logistic regressions). Correlations are with the first member of the pair listed, with the second member of the pair as the index value. Body mass is log-transformed; plants and vertebrates are dietary categories scored, along with invertebrates, out of a total of 100; mating system coefficients are against a baseline of monogamy; all continuous variables are scaled to have a mean of 0 and a variance of 1 prior to analysis. Significant correlations are highlighted in grey (pMCMC < 0.05).

Both male- and female-biased SSD, however, are correlated with mating system. Compared with monogamous species, male-biased SSD is more common in polygynous (β_m_ = 1.324, pMCMC = 0.002) and polygynandrous (β_m_ = 1.002, pMCMC = 0.034) species, whilst female-biased SSD is much more common in polyandrous species (β_f_ = 3.049, pMCMC = 0.014). There is also an increase in male-biased SSD within the “mixed” mating system category in the separate effects model (β_m_ = 1.058, pMCMC = 0.030, Table 1) that is not recapitulated in the logistic regressions (Table 2).

These results are generally robust to both the threshold used to determine SSD and to scores based on linear measurements instead of body mass, with some minor variations in the p-value estimate of the seasonality variables (see Tables S7-9).

## Discussion

Across all mammals, sexual monomorphism is a relatively stable state, and female-biased SSD is a relatively unstable state. Transitions between these states are correlated primarily with variation in mating system and body mass, with a smaller role of diet and seasonality, and no statistically meaningful correlation with social organisation. While male-biased SSD is more common in larger species, after accounting for variation due to mating system there is no difference in body mass between species with female-biased SSD and monomorphic species; thus, Rensch’s rule is only partially supported at the class-wide scale.

### Patterns of SSD state transition

The analysis of macroevolutionary transitions in SSD suggest that monomorphism is an evolutionarily stable state among mammals, and thus that if the selective pressure acting separately on male and female body size is subsequently removed, a lineage is likely to quickly lose any dimorphism. In other words, these results suggest that dimorphism may be – on a macroevolutionary timescale – costly to maintain, especially female-biased SSD. This aligns with the observation that female-biased SSD is far less common among mammals than male-biased SSD is [1, 2]. Furthermore, this may suggest that the predominance of male-biased SSD among mammals is partially due to these sex-based differences in evolutionary rates from SSD back to monomorphism, if selection acts more strongly to return female body size to the species-wide average than it does to male body size. However, the observed results could also be caused by other factors: for example, if female-biased SSD were, on a geological timescale, a much older trait than male-biased SSD, there may have been less time in which male-biased SSD could be secondarily lost. Though this particular situation seems unlikely, given evidence from the fossil record and ancestral state reconstructions [17, 40, 41], further work would need to be done to determine exactly why male-biased SSD appears to be more evolutionarily stable, and what this means for our understanding of sex-specific body size evolution.

### Rensch’s rule

Though we find strong evidence that male-biased SSD is more common in larger animals (in accordance with Rensch’s rule), we also find that female-biased SSD is positively correlated with mass, and that there is no difference in body mass between species with female-biased SSD and monomorphic species (in disagreement with Rensch’s rule, which predicts that smaller species should have higher female-biased SSD) [12, 14, 16]. Thus, our results suggest that Rensch’s rule is not fully applicable at a whole-Mammalia level, a finding which accords with many comparative studies (e.g. [3, 12, 16]). This could in part be due to our multivariate modelling approach, which controls for factors such as mating system that are often posited as the mechanical link underlying the pattern Rensch’s rule describes [12, 13]. Alternatively, it could be that fecundity selection – the hypothesis sometimes suggested to explain Rensch’s rule, that larger species have more constrained female body masses because reproduction is more costly in these groups [3] – creates a greater difference in optimal male and female body size [18], but does not necessarily result in selection pressure for smaller females.

### Seasonality and SSD

Temperature seasonality is the least well-researched of the variables which we found to be significantly correlated with SSD, and the few existing studies which investigate the link between SSD and seasonality are frequently contradictory [4]. Furthermore, the relationships between seasonality and SSD established here were the only results found to be sensitive to reassignments in the dataset according to dimorphism threshold (Tables S7-8) or morphological proxy (Table S9); and any effects of terrestrial seasonality will be less applicable to some ecological strategies, such as aquatic or migratory species. Thus, interpreting this correlation is challenging. A common method of explaining this link appeals to Rensch’s rule, as species at highly seasonal environments tend also to be larger [4]; it is also possible that an SSD-seasonality correlation is driven by an underlying relationship between both variables and social organisation [28]. Our models, however, find a correlation between seasonality and male-biased SSD above and beyond the correlations with body size and sociality (see Table S2), and the overall statistical support for the relationship with social organisation is low. Furthermore, the low VIFs indicate that any correlations between these variables are not distorting our model results and are not confounding this SSD-seasonality relationship.

Prior studies on prosimians, particularly those from Madagascar, have suggested that increased seasonality correlates with reduced SSD in this group [25, 26], in contrast to our findings. Garel *et al*. [27], however, found that Norwegian moose populations in more seasonal regions show greater male-biased SSD. The authors attribute this to shorter growing seasons producing more easily digestible plants, and males being more heavily selected to prioritise growth than females, resulting in males seeing a greater increase in body mass when better-quality forage is available. However, while the patterns observed in this study align with our whole-Mammalia results, this plant-foraging explanation is difficult to apply across the diets of all mammals. It is unlikely that increased seasonality increases forage quality worldwide, given that the tropics have much greater biomass productivity than temperate regions [42].

Another potential explanation for the correlation of temperature seasonality and male-biased SSD is that seasonality facilitates the initial evolution of SSD via seasonal dimorphism. Ferretti *et al*. [20] suggest that seasonally dimorphic species such as the Apennine chamois represent an evolutionary midpoint between monomorphic and permanently dimorphic species. If this were the case, the link between temperature variability and SSD may reflect an underlying link between both traits and seasonal dimorphism. The evidence to support this hypothesis at the class-wide scale, however, is currently lacking; the incorporation of other types of dimorphism into future studies of SSD may yield clarifying results.

### Diet and SSD

The association of vertebrate predation with male-biased SSD supports prior observations among the Carnivora – particularly the mustelids and their close relatives – that species with a greater reliance on vertebrate prey tend to have more pronounced male-biased SSD [17, 21, 22]. The main hypothesis proposed to explain this pattern is that the uneven spatial distribution of vertebrate prey promotes intra-specific competition for food, more so than plants or invertebrate prey do [17, 21]. Thus, there is a greater selection pressure for reducing intra-specific competition. Niche divergence between the sexes, mediated by sexual differences in body size, is one method to achieve this [17, 21]. However, this hypothesis does not predict that one sex in particular would become larger than another, merely that some sort of SSD will develop. If this was the only factor driving the association of carnivory with SSD, both female-biased and male-biased SSD would be expected to have an association with vertebrate predation. Instead, this correlation only applies to male-biased SSD, suggesting that this link is one of several evolutionary pressures shaping the present-day distribution of this trait.

Noonan *et al*. [22] link carnivory back to the sexual selection hypothesis, suggesting that the more clustered distribution of vertebrate prey promotes spatial clustering in the predatory species, which in turn facilitates female monopolisation by males (polygyny). This link has the potential to explain why only male-biased dimorphism is positively associated with vertebrate predation in our results. However, both Noonan et al.’s study and ours only test for correlations, not evolutionary pathways, so cannot indicate any particular chain of causation. We note, however, that low VIF values in our mammal-wide models suggest that vertebrate predation and polygyny are not highly correlated in our sample.

Another possible explanation for carnivory driving male-but not female-biased SSD is that factors such as fecundity selection may limit female body mass [3, 17]. In this case, dietary niche divergence would promote an increase in male, but not female, body size, leading to a predominance of male-biased SSD among species which experience high degrees of intraspecific competition for food.

Alternatively, the lack of correlation between female-biased SSD and vertebrate predation may be biased by the importance of the Carnivora to this correlation. The Carnivora – the group which has previously been the focus of studies linking vertebrate predation to male-biased SSD – generally have a far greater percentage of vertebrates in their diet than any other clade of mammals (Figure 2). They also primarily display male-biased SSD, although female-biased SSD occurs rarely. If this particular group is more disposed towards male-biased SSD, for example due to genetic features in their common ancestor, and they are also the only group with a large degree of vertebrate predation, then this may explain the lack of correlation between female-biased SSD and vertebrate predation, without requiring any reasons why vertebrate predation and/or niche divergence would inherently favour larger males.

Further studies on the link between SSD and dietary niche divergence will be complicated by the lack of global, detailed mammalian dietary data [23, 24]. Furthermore, SSD may plausibly be a cause, rather than an effect, of niche divergence [1], and correlation and causation would be difficult to disentangle. Our results, however, do support diet, and potentially dietary niche divergence, as an important contributor to the broad-scale evolution of SSD in modern mammals.

### Mating Systems and SSD

Our results linking SSD to mating systems align with previous studies and observations at a variety of scales (e.g., [3, 5, 43-45]). We find, for example, that polyandry is associated with female-biased SSD, while polygyny is associated with male-biased SSD.

The most popular and persistent explanation in the literature for the correlation between mating systems and SSD involves pre-copulatory sexual selection. Larger males are proposed to have a competitive advantage in direct, aggressive competition for females and/or to be preferred mating partners of females. Among polygynous species, this would mean that larger males would mate with more females, producing more offspring, and so sexual selection would favour large size among males, which will eventually lead to the evolution of male-biased SSD [1, 2, 4, 5]. The converse might plausibly apply to polyandrous species and female-biased SSD, particularly in group-living species [10, 11]. By contrast, in monogamous species, each individual will have at most a single mate, reducing the benefit any competitive advantage due to size can convey. Our results generally align with this hypothesis, although the slight positive association between polyandry and male-biased SSD recovered by our logistic regressions would indicate a more nuanced relationship between mating systems and SSD. Future studies across a larger sample of species, encompassing a greater evolutionary and ecological context, together with more detailed social organisation data and group size data, may help elucidate the complicated relationship between species-level social behaviour and sex-specific body size evolution.

Recent genetic paternity studies, however, have thrown doubt on the validity of this sexual selection hypothesis [5, 8, 9]; in part, it seems that many species typically considered polygynous are in fact polygynandrous. Moreover, in order for sexual selection in polygynous species to lead to male-biased SSD, a population must a) show variation in male reproductive success, and b) have this variation significantly correlated with male body size, with larger males achieving greater reproductive success [46], but these criteria are not always met in mammals. For example, no reliable correlation between SSD and variation in male reproductive success has been found in studies of pinnipeds and primates [7, 8]. Cassini [47] did find that SSD correlated with intensity of sexual selection among artiodactyls, but it seems clear that this pattern is not universal throughout Mammalia. Furthermore, the evolution of SSD appears to predate the evolution of polygyny in pinnipeds (the mammalian clade with the greatest degree and variety of SSD), suggesting that simple correlation would not adequately support the clear cause-and-effect pathway suggested by sexual selection [48].

Cassini [49] found that the pattern of SSD evolution in artiodactyls could best be explained through a model where an increase in body mass, diet specificity, and sociality – all driven by a move to a more open environment – led to the evolution of polygyny, which in turn promoted sexual segregation, which then drove the evolution of SSD. This particular pattern of causation is highly unlikely to apply across Mammalia – e.g., Krüger *et al*. [48]’s finding that polygyny evolved after SSD in pinnipeds would preclude polygyny from triggering the evolution of SSD in this group. However, Cassini’s study does offer an example of how a polygynous mating system could lead to the evolution of SSD without sexual selection being the main driving force, and also of how all of the factors analysed in this study – along with a few others – may interact to drive SSD evolution.

## Conclusion

Through modelling macroevolutionary transition rates, we found that SSD is more likely to be lost than gained, and thus is an evolutionarily unstable trait. Both female-biased and male-biased SSD evolved from monomorphism at similar rates, but female-biased SSD was lost at a greater rate, potentially leading to the predominance of male-biased SSD seen in today’s mammals. Further study, incorporating the evolutionary history of these traits, could illuminate whether these results do indeed imply that female-biased SSD is inherently costly to maintain on an evolutionary timescale.

Body mass, mating system, diet, and seasonality are all significantly correlated with presence of SSD across the mammalian class, though any link with social organisation was considered tenuous at best. Most of these broad-scale effects were in line with previous observations and hypotheses at smaller scales, with larger, polygynous, and/or carnivorous species more likely to display male-biased SSD, and with polyandrous species more likely to display female-biased SSD. However, contrary to the predictions of Rensch’s rule, large size also weakly favoured female-biased SSD. We also particularly highlight the relationship of seasonality with SSD as a potential link for further study, particularly as it may or may not relate to social organisation. Overall, our models suggest that many social and ecological factors work together to drive and maintain broad-scale evolutionary patterns of mammalian sexual size dimorphism.

## Author Contributions

MEJ analysed the data and wrote the first draft of the manuscript. CS designed the study and collected the data. Both authors contributed to the writing and editing of the manuscript.

## Acknowledgments

This project was funded by ERC grant 788203 (Innovation). We thank Michael Benton and the Bristol Macroevolution lab group for early project feedback as well as Chris Law and two anonymous reviewers for thoughtful, constructive comments on a previous version of this manuscript.

